# A microfluidic platform with integrated porous membrane cell-substrate impedance spectroscopy (PM-ECIS) for biological barrier assessment

**DOI:** 10.1101/2023.11.25.568615

**Authors:** Alisa Ugodnikov, Joy Lu, Oleg Chebotarev, Craig A. Simmons

## Abstract

Traditionally, biological barriers are assessed in vitro by measuring trans-endothelial/epithelial electrical resistance (TEER) across a monolayer using handheld chopstick electrodes. Implementation of TEER into organ-on-chip (OOC) setups is a challenge however, due to non-uniform current distribution and interference from biomaterials typically found in such systems. In this work, we address the pitfalls of standard TEER measurement through the application of porous membrane electrical cell-substrate impedance sensing (PM-ECIS) to an OOC setup. Gold leaf electrodes (working electrode diameters = 250, 500, 750 µm) were incorporated onto porous membranes and combined with biocompatible tape to assemble microfluidic devices. PM-ECIS resistance at 4 kHz was not influenced by presence of collagen hydrogel in bottom channels, compared to TEER measurements in same devices, which showed a difference of 1723 ± 381.8 Ω (p=0.006) between control and hydrogel conditions. A proof of concept, multi-day co-culture model of the blood-brain barrier was also demonstrated in these devices. PM-ECIS measurements were robust to fluid shear (5 dyn/cm^2^) in cell-free devices, yet were highly sensitive to flow-induced changes in an endothelial barrier model. Initiation of perfusion (0.06 dyn/cm^2^) in HUVEC-seeded devices corresponded to significant decreases in impedance at 40 kHz (p<0.01 for 750 and 500 µm electrodes) and resistance at 4 kHz (p<0.05 for all electrode sizes) relative to static control cultures, with minimum values reached at 6.5 to 9.5 hours after induction of flow. Our microfluidic PM-ECIS platform enables sensitive, non-invasive, real-time measurements of barrier function in setups integrating critical OOC features like 3D co-culture, biomaterials and shear stress.

## 1.0 Introduction

Biological barriers play a critical role in maintaining homeostasis, combining tissue-specific endothelial or epithelial cells, matrices, support cells, junctions and transporters to regulate the movement of nutrients and pathogens between compartments of the body and the external environment^1–3^. The gold standard techniques for assessing biological barriers *in vitro* are the molecular tracer permeability assay and trans-epithelial/endothelial electrical resistance (TEER). While the permeability assay is amenable to microfluidic setups, TEER is more challenging to implement. Traditionally, TEER is measured across a monolayer grown on a cell culture insert using handheld chopstick electrodes. Background resistance of the system is accounted for by measuring blank inserts and subtracting the corresponding resistance from cell-seeded inserts, and the resulting value is multiplied by the exposed membrane area to allow comparison across various insert sizes. These calculations, however, do not account for the non-linear fashion in which the current path increases as the size of the cell culture insert increases^4^. Non-homogenous current density distribution, particularly pronounced with chopstick electrodes^5^, results in a smaller measurement area relative to total membrane area as insert size increases^4^, yielding significantly different TEER values for the same cell types^6^. In microfluidic devices it is especially difficult to ensure a uniform current distribution with electrodes placed in channel outlets, due to low conductivity within microfluidic channels^7^. Integrating the electrodes directly into the top and bottom channels addresses this limitation, but is challenged by the small device dimensions and constraints for cell imaging, which restrict the parameters that determine electric field uniformity, e.g., electrode size and placement^7,8^. Although efforts have been made to create correction factors to help translate between microfluidic and Transwell TEER values^4,5,7^, conditions specific to organ-on-chip culture may introduce additional error^9^.

Since TEER electrodes, whether inserted into channel outlets or integrated into the device, are located at a distance from the cell layer, they are measuring not only the cells but rather the bulk impedance of all the materials between the electrodes. This is particularly relevant to OOC platforms as many incorporate hydrogels that can limit the sensitivity of TEER to changes in the cell layer of interest. Hydrogels commonly used in OOC are high viscosity, which contributes to increased resistances, and have low inherent conductivity^6,10^. For example collagen, collagen-hyaluronic acid, and Matrigel hydrogels have reported conductivities in the range of 0.02 – 0.6 mS/cm^11–13^, which is 2 to 3 orders of magnitude lower than that of cell culture medium^14,15^. Additionally, the presence of cells^10^, cellular phenomena - like gel contraction and deposition or degradation of extracellular matrix^16^ - or even irregularities such as air bubbles^17,18^ can all influence the electrical properties of a hydrogel. In such culture systems, with high inherent resistance and where multiple components are interacting over time, interpreting data acquired through electrodes in the standard parallel configuration can be challenging^16^.

Electrical cell-substrate impedance sensing (ECIS), where cells are grown directly on planar electrodes, provides an alternative method for sensitive measurements of monolayers^19–21^. Alternating current is applied across the cell layer over a range of frequencies, which together with an equivalent circuit model can be used to provide information about phenomena such as cell proliferation, migration and barrier integrity; the theory behind ECIS is reviewed elsewhere^22–25^. The classic setup uses a single small, circular (d=250 µm) sensing electrode and a large counterelectrode integrated onto the bottom of a well plate, though various configurations are available from Applied Biophysics, including interdigitated electrodes and flow arrays for studying the effect of fluid shear^26^. Beyond commercially available products, research groups have developed customized microfluidic ECIS systems to evaluate biological barriers under flow. Lewis *et al.* incorporated optically transparent, interdigitated indium tin oxide electrodes to allow for simultaneous visualization of HUVEC monolayers^27^; more recently, Velasco *et al.* tested three different electrode sizes (d = 50, 100, 200 µm) for their sensitivity to detect flow-induced changes in HUVECs^28^. The majority of ECIS platforms, however, incorporate the electrodes onto a solid substrate of glass or thermoplastic. This presents an obstacle to evaluating co-cultures, as well as validating ECIS measurements against standard barrier assessment techniques, which require a porous membrane substrate^29^.

Recent efforts to address this gap – Rajasekaran *et al.* in Transwell inserts^30^; Schuller *et al.* and Matthiesen *et al.* in microfluidic devices^31,32^ – use e-beam deposition or sputtering to incorporate gold electrodes directly onto porous membrane. Our previous work^33,34^ has demonstrated that porous membrane ECIS (PM-ECIS) electrodes fabricated using lower cost gold leaf are sensitive to changes in endothelial barriers in a cell culture insert embodiment, and that these measurements can be simultaneously validated against standard TEER and molecular permeability barrier assessment methods. Here we present the implementation of PM-ECIS in a cost-effective bilayer microfluidic barrier-on-chip platform fabricated from acrylic and double-sided adhesive. We show that PM-ECIS measurements in this platform are robust to the presence of hydrogel in bottom channels and fluid flow-induced shear stress, can be made over multiple days in co-culture and perfusion culture, and can detect flow-induced changes in endothelial barrier integrity with electrode-size dependent sensitivity. These data further the understanding of ECIS on porous membranes and support the potential of PM-ECIS as a technology that enables comparison of biological barrier assessment across various *in vitro* platforms.

## 2.0 Materials and methods

### 2.1 Cell culture

GFP-expressing human umbilical vein endothelial cells (HUVECs) (AngioProteomie, cAP-0001GFP) were thawed and suspended in Endothelial Growth Medium (AngioProteomie, cAP-02). T75 flasks were pre-coated with 4 mL Quick Coating Solution (AngioProteomie, cAP-01) at room temperature for 5 min, then aspirated. HUVEC cell suspension (1−10^6^ cells) was transferred into the flask and placed overnight into a cell culture incubator (37°C, 5% CO_2_) to allow cells to adhere. Medium was replaced with 15 mL fresh EGM the next day, and changed every other day thereafter. HUVECs were cultured for 6-7 days prior to detachment with 0.25% trypsin-EDTA (Life Technologies, 25200-056) for use in experiments.

Immortalized human brain microvascular endothelial cells (HBMEC) (provided by Tara Moriarty, University of Toronto)^35^, were cultured using the EGM™-2 MV Microvascular Endothelial Cell Growth Medium BulletKit™ (Lonza, CC-3202), which contains hydrocortisone, and was supplemented with 20 µg/mL Hygromycin B Biobasic Canada (Cat #BS725) to select for transfected cells. Primary human astrocytes (Thermo Fisher, N7805100) were used for the co-culture conditions and grown in DMEM supplemented with 10% FBS, 1% penicillin-streptomycin and N-2 supplement (17502-048).

### 2.2 Device fabrication

Electrodes were incorporated onto a porous transparent PETE membrane (3.0 µm, 6−10^5^ pores/cm^2^; Sterlitech, 1300032) using thermal bonding of gold leaf (Gold Leaf Supplies, 2300/RUXX) to the membrane, followed by UV lithography to pattern the electrodes and laser cutting of the electrode-patterned membrane to size and shape, using the protocol previously described^33,34^. An insulating mask (Grand & Toy, #99842) was used to cover electrical traces, and the exposed gold served as electrical contacts to connect via a flexible printed circuit board to a 24-channel multiplexer, lock-in amplifier (LIA)) and computer, connected as in our previously described setup^33,34^.

PM-ECIS electrodes were incorporated with layers of acrylic and bio-compatible double-sided tape (ARSeal 90880, Adhesives Research) to form microfluidic channels containing three working electrodes (diameters = 250, 500 and 750 µm) (**Figure 1**). The spacing of the counter electrode (area = 0.075 cm^2^) with respect to each working electrode was 900 µm, measured by taking the shortest edge-to-edge distance between each working electrode and the counter electrode. The top and bottom channels, of equal dimensions (width = 3 mm, height = 284 µm), were generated by laser cutting a channel pattern into a sheet formed by laminating two layers of the double-sided tape. Inlets and outlets were laser-cut in an acrylic top layer to allow for insertion of tubing for connection to the syringe pump. A single layer of the tape was placed between the acrylic and top channel to facilitate a tight seal. HumiSeal UV40 (Chase Corporation), a biocompatible UV curable conformal coating, was then applied to the outer walls of the devices to prevent leakage and UV cured following product guidelines^36^.

**Figure 1:**
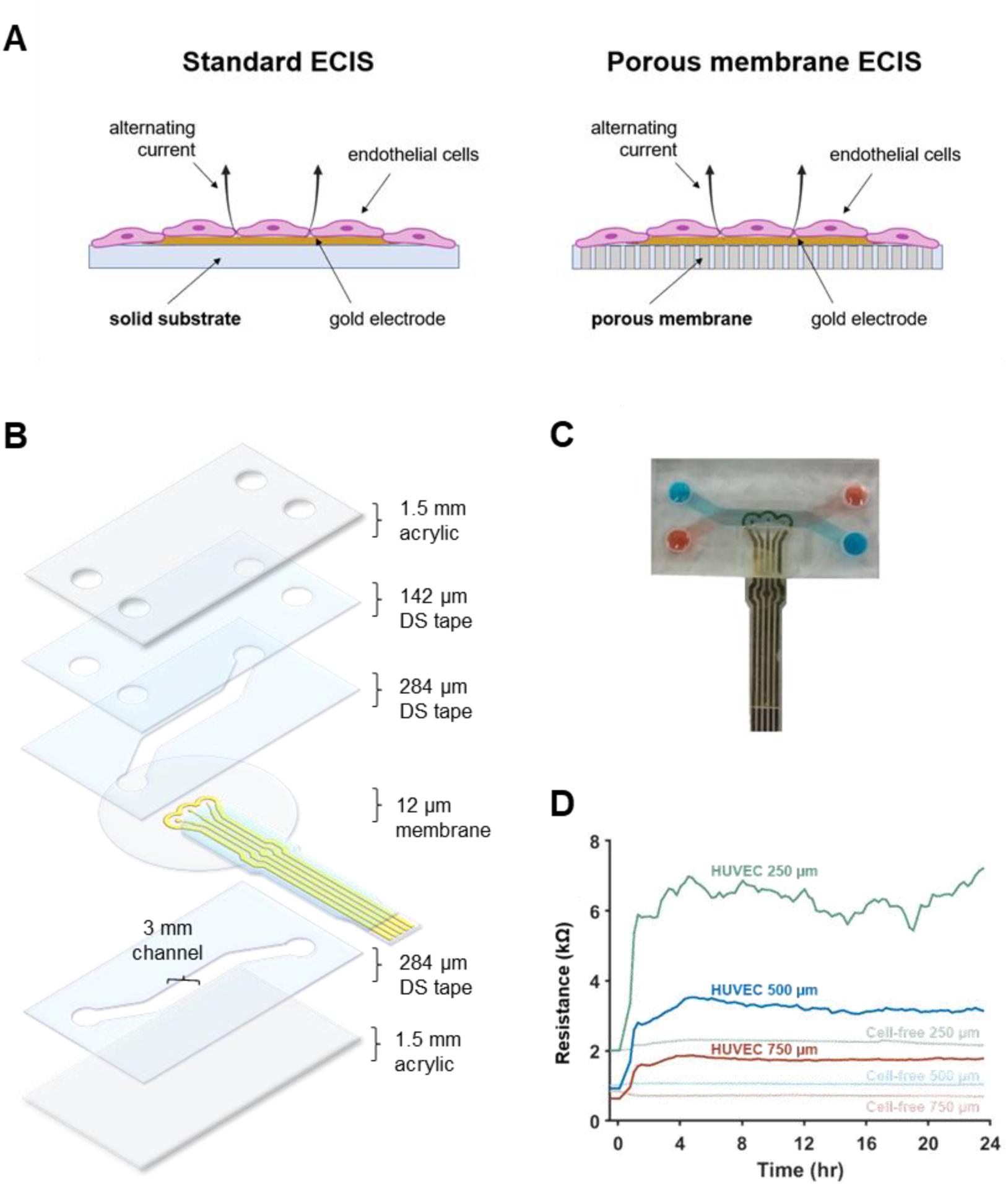
Microfluidic ECIS platform. **A**: Schematic diagram comparing standard ECIS vs. PM-ECIS electrodes, electrode thickness not to scale. Schematic created using BioRender. **B:** Fabrication of microfluidic device incorporating porous membrane (PM-ECIS) electrodes. **C:** Top-down view of device with integrated PM-ECIS) electrodes. **D:** Resistance at 4 kHz for HUVECs seeded at t=0 on working electrodes of various sizes (d = 250, 500, 750 μm). Resistance values for corresponding cell-free control electrodes shown for comparison. DS tape = double-sided tape.

### 2.3 ECIS and TEER measurements

ECIS measurements were performed using methods described previously^33,34^. A cell-free baseline was established beforehand using cell-free devices filled with culture medium and impedance measurements taken over the course of at least 12 hours prior to cell seeding. LIA outputs including voltage drop and phase angle were used to calculate and plot impedance and resistance values using a custom MATLAB program^33^. Measurements for cell-based experiments reported for 4 kHz and 40 kHz. Frequency scans involved measurements over 11 frequencies (62.5, 125, 250, 500, 1000, 2000, 4000, 8000, 16000, 32000, 64000 Hz).

TEER was measured using a Millicell ERS volt-ohm meter (World Precision Instruments, New Haven, CT). 200 µL reservoirs filled with culture medium were inserted into the channel outlets and electrodes placed in reservoirs to take measurements; total exposed membrane area was 1.9 cm^2^.

### 2.4 Microfluidic device culture

Devices were sterilized by filling top and bottom channels with 70% ethanol for 30 min at room temperature, then washed with PBS and UV sterilized for 1 hour. Top channels (exposed membrane area ∼ 1.05 cm^2^) were filled with 30 µL of 50 µg/mL bovine plasma fibronectin (Sigma-Aldrich, F1141) at for 90 min at room temperature, then both top and bottom channels filled with culture medium and connected to the ECIS system to allow for stabilization of measurements for at least 12 h.

To test effects of hydrogels on ECIS and TEER measurements, bovine collagen type I (2.7 mg/mL) gel containing DMEM was incorporated in bottom channels in some cases. 10X DMEM solution was made by combining 2.7 g powdered DMEM (Sigma-Aldrich, D7777-10L) and 0.74 g NaHCO_3_ in distilled water in a total volume of 20 mL, pH = 7.4. To make the collagen hydrogels, 125 µL 10X DMEM solution, 968 µL collagen (PureCol bovine collagen type I, Advanced Biomatrix, 5005-B) and 15 µL 0.8M NaHCO_3_ were combined on ice. All control devices contained DMEM culture medium only in both top and bottom channels. First, baseline ECIS measurements were taken for all devices. Then, for the devices in the collagen condition, 50 µL of the collagen solution was added to the bottom channel and devices placed in an incubator at 37°C for 30 min to allow polymerization. Subsequently, ECIS measurements were taken again for all devices 2 hours after the polymerization was completed (“2 h post-polymerization”) in the collagen condition devices. Change in resistance values at 4 kHz were calculated using Δ R = R_2h post-polymerization_ – R_baseline_. For ECIS vs. TEER comparison, measurements of medium-only control devices and collagen condition devices were taken after 1 day of incubation at 37°C, 5% CO_2_,

For experiments with HUVECs, cells (P5) were suspended in EGM at a density of 3−10^6^ cells/mL and 30 µL was injected into the top channel, where the cells were allowed to attach overnight. Medium was changed daily thereafter. For blood-brain barrier cultures, immortalized HBMEC were seeded in top channels (30 µL of 3−10^6^ cells/mL cell suspension) and allowed to attach for 1 hour at 37°C. Astrocytes were gently mixed in collagen type I solution containing DMEM (prepared as described above) at 1−10^6^ cells/mL. 30 µL of the astrocyte/collagen suspension was added to the bottom channels, and devices incubated at 37°C for 1 hour to polymerize the collagen gel, with the devices upside to ensure proximity of astrocytes to the porous membrane. Endothelial medium was used for media changes, performed daily.

### 2.5 Application of fluid flow

Devices were connected to a syringe pump for application of fluid flow. For assessing the effect of flow in cell-free devices, channels were filled with phosphate-buffered saline (PBS) without Mg^2+^/Ca^2+^, and measurements taken at room temperature before and after application of fluid flow-induced wall shear stress (5 dyn/cm^2^). Change in resistance was calculated by subtracting ECIS resistance values immediately prior to induction of flow from ECIS resistance values immediately following (Δ R = R_post-flow_ – R_pre-flow_). For HUVEC culture, medium was degassed using vacuum filtration, and a flow rate of 20 uL/min (equivalent to a wall shear stress of 0.06 dyn/cm^2^) applied 2 days after cell seeding, continued for up to 5 days of flow. The minimum impedance (40 kHz) and resistance (4 kHz) values after induction of flow (“post-flow minimum”) were obtained for flow condition electrodes, and compared to static condition at the same respective time points for each electrode size.

### 2.6 Immunostaining

Cells were fixed in 10% neutral buffered formalin for 10 min at room temperature, washed three times with PBS containing Mg^2+^/Ca^2+^, permeabilized with 0.2% Triton-X in PBS, and blocked with 3% bovine serum albumin (BSA) (Sigma-Aldrich, 10735086001) for 20 min at 37°C. Cells were stained with primary antibodies (1:100) diluted in 3% bovine serum albumin in PBS-T (0.1% Triton-X in PBS) for 1 hour at 37°C. Following incubation, cells were washed three times with PBS and blocked using 10% goat serum (Sigma-Aldrich, G9023) in PBS-T for 1 hour at room temperature. Secondary antibodies (1:200) diluted in 10% goat serum were applied for 1 hour at room temperature. After washing three times with PBS, nuclei were stained with 1:1000 Hoechst 33342 (Sigma-Aldrich, B2261) for 5 min at room temperature. Primary antibodies were rabbit polyclonal vascular endothelial (VE)-cadherin (Abcam, ab33168), mouse monoclonal zonula occludens-1 (ZO-1) (Thermo Fisher Scientific, 33-9100), rabbit polyclonal glial fibrillary acidic protein (GFAP) (Abcam, ab7260); secondary antibodies were goat anti-mouse IgG (H+L) Alexa Fluor 647 (Life Technologies, A-21236) and goat-anti rabbit IgG (H+L) Alexa Fluor 568 (Thermo Fisher Scientific, A-11011). Image acquisition was performed using an Olympus FV3000 confocal laser scanning microscope or Olympus BX51 inverted fluorescence microscope as indicated.

### 2.7 Statistical analysis

Statistical analysis was performed using GraphPad Prism 5. Unpaired t-test was used to compare collagen and control condition resistance values for chopstick TEER *vs.* PM-ECIS in cell-free devices. One-way ANOVA with Bonferroni test was used to analyze collagen and control condition measurements across PM-ECIS electrode sizes. A one sample t-test with theoretical mean of 0 was used to assess if induction of fluid flow in cell-free devices influenced PM-ECIS resistance measurements at 4 kHz. For HUVEC-seeded devices, an unpaired t-test was used to compare PM-ECIS measurements before and after application of flow, and one-way ANOVA with Bonferroni test was used to assess the effect of electrode size on post-flow minimum measurements. Unpaired t-test was used for each of the first 6 timepoints after induction of perfusion to compare static vs. flow conditions. Samples (N) represent independent devices.

## 3.0 Results

### 3.1 Effect of collagen hydrogel on ECIS resistance vs. chopstick TEER

Organ-on-a-chip configurations of membrane-based microfluidic systems often include a hydrogel in the abluminal channel, the presence of which alone could affect cell barrier impedance measurements. To test this, the resistances of cell-free membranes were measured by ECIS and chopstick TEER in devices containing DMEM in the top channel and either DMEM (control) or collagen hydrogel (collagen) in the bottom channel (**Figure 2A**). Importantly, presence of hydrogel alone in the bottom channel did not influence ECIS resistance, compared to TEER measurements in same devices, which showed a difference of 1723 ± 381.8 Ω (p=0.006) between control and hydrogel conditions (**Figure 2B**). In contrast, the resistance measurements performed by the ECIS electrodes were robust to the presence or absence of hydrogel in the bottom channel across all electrode sizes, with no significant difference (p=0.44) between the control and hydrogel conditions (**Figure 2C**).

**Figure 2:**
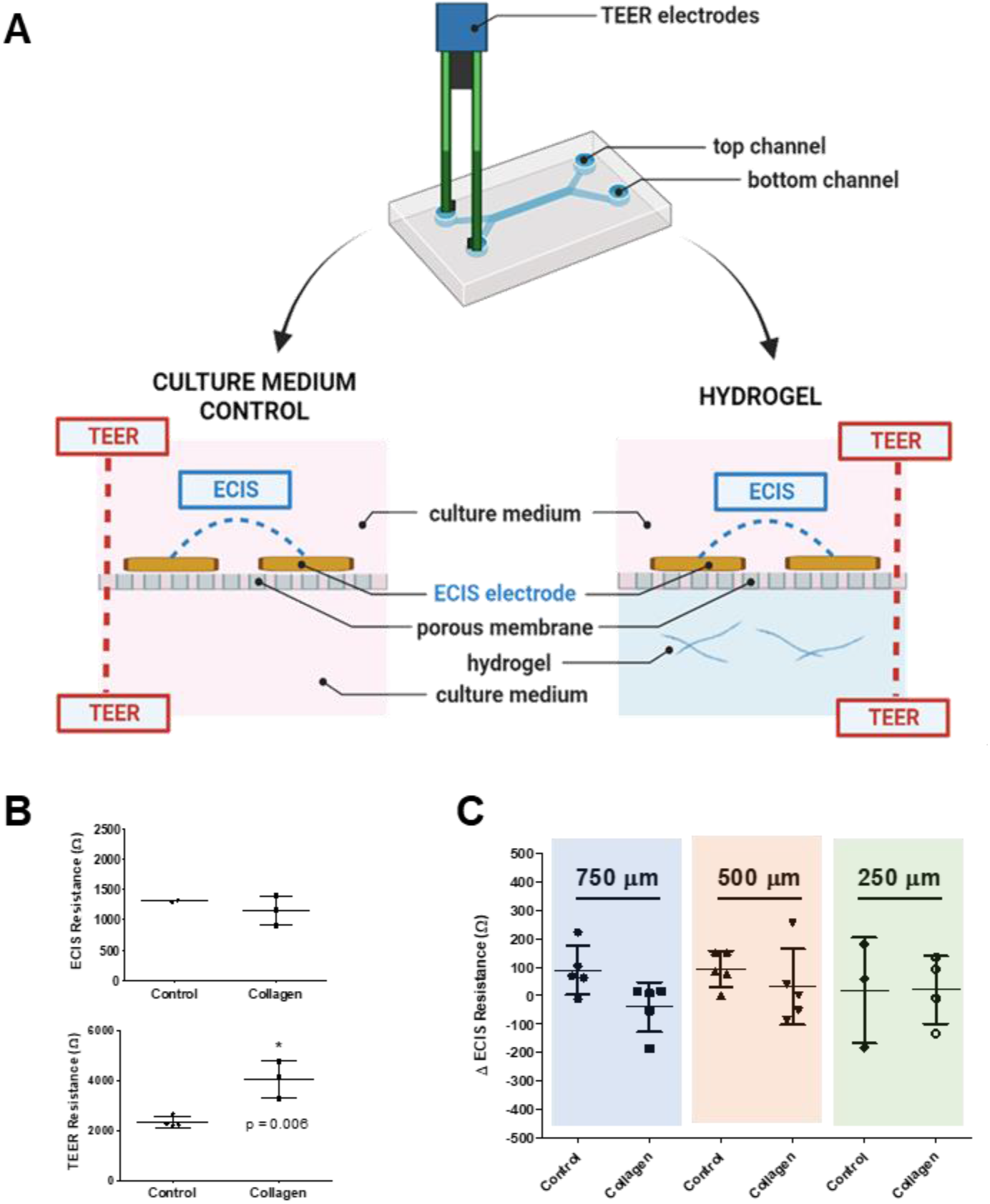
Effect of collagen hydrogel on ECIS resistance vs. chopstick TEER. **A**: Measurement setup for ECIS resistance and chopstick TEER. Schematic created using BioRender **B:** Resistance values at 4 kHz for devices with collagen gel in bottom channel vs. control for ECIS and TEER, respectively, at Day 1, N= 2-4 per condition. Unpaired t-test, *\* p < 0.01.* **C:** Change in resistance values at 4 kHz for devices with collagen gel in bottom channel vs. medium-only control at 2 hours post-polymerization (t=2.5 h) for various working electrode diameters (Δ R = R_2h post-polymerization_ – R_baseline_); N=3-5 per condition. No significant difference between control and collagen conditions (p=0.44 by one-way ANOVA). Data presented as mean ± standard deviation.

### 3.2 Blood-brain barrier model in microfluidic ECIS platform

Having established that ECIS resistance measurements in the microfluidic platform were unaffected by the presence of a hydrogel, we tested the platform in a relevant 3D co-culture model of the blood-brain barrier. HBMECs were grown on the luminal side of the porous membranes with the ECIS electrodes, and primary human astrocytes embedded in collagen hydrogel were grown in the abluminal channel (**Figure 3A**) for up to 5 days. HBMECs were evident on the porous membrane and electrodes on the luminal side by their expression of the tight junction zonula occludens-1 (ZO-1) (**Figure 3B,C**), while astrocytes expressing glial fibrillary acidic protein (GFAP) were evident on the abluminal membrane surface and spanned the membrane-electrode border (**Figure 3D,E**). The platform supported BBB co-culture over the course of 5 days, reaching peak barrier integrity on Day 3 post-seeding as measured by ECIS resistance at 4 kHz using the 500 µm diameter electrodes (**Figure 3F**).

**Figure 3:**
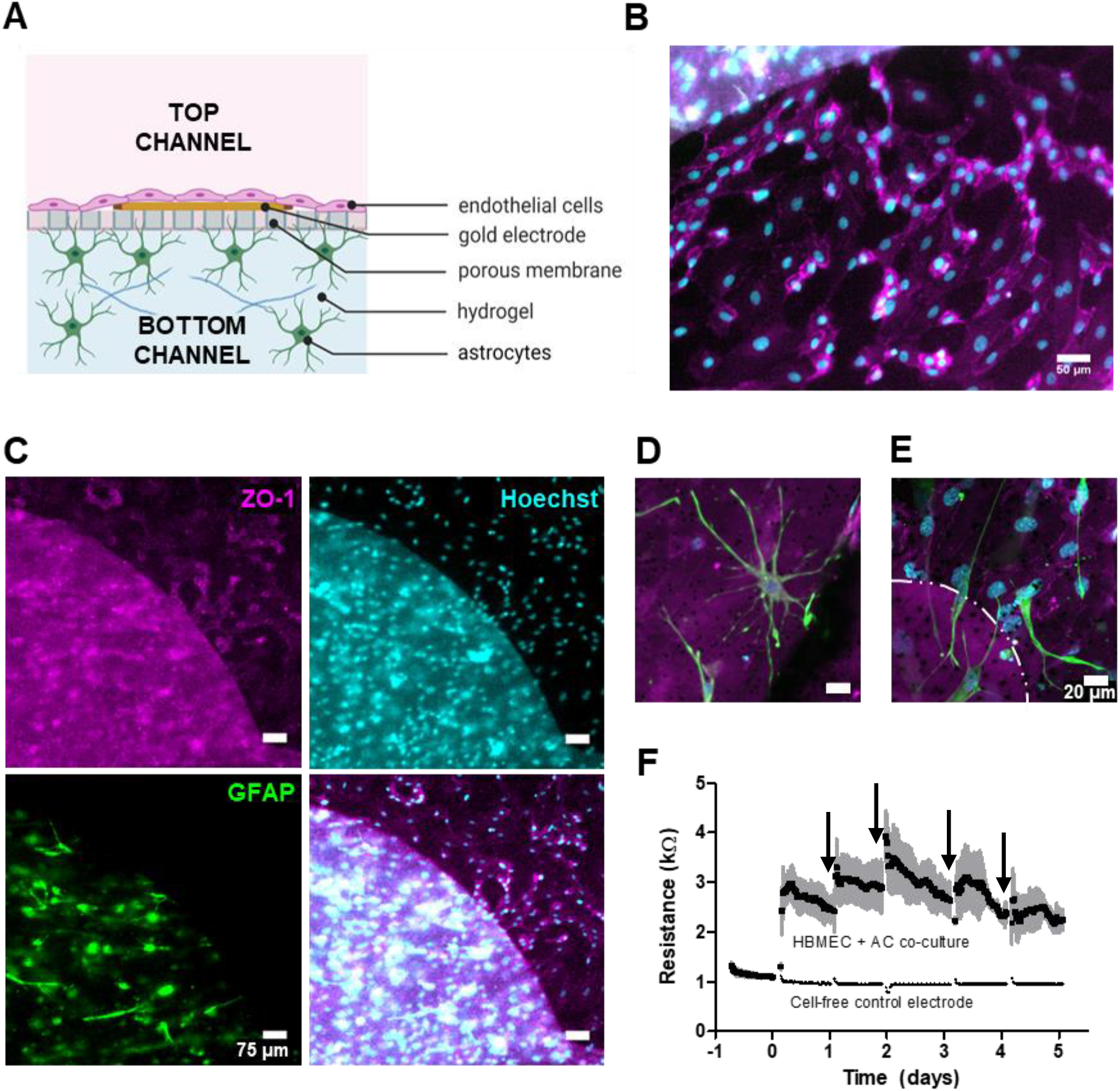
Blood-brain barrier model in microfluidic ECIS platform. **A:** Schematic of blood-brain barrier (BBB) co-culture setup, with HBMEC in top channel and primary human astrocytes (AC) in bottom channel. Schematic created using BioRender. **B, C:** Fluorescence microscopy imaging from top side, showing HBMEC cell layer on counter electrode surface (black), and AC embedded in hydrogel underneath**. D, E:** Confocal imaging of BBB co-culture in microfluidic setup from astrocyte side of membrane. Dotted line indicates border between working electrode and membrane. AC stained for GFAP (green), HBMEC stained for ZO-1 (pink) and nuclei visualized with Hoechst 33342. **F:** ECIS measurements of BBB co-culture over 5 days in static condition, showing traces for three different co-culture devices. Cell seeding occurred at t=0; black arrows = media changes. Mean ± SD plotted for N=3 independent electrodes (d=500 µm) for co-culture condition.

### 3.3 Microfluidic ECIS measurements upon application of fluid flow

A significant advantage of microfluidic ECIS is the ability to perfuse cells in the luminal channel. To first validate that ECIS measurements were not affected by fluid flowing over the electrodes on the porous membrane, PBS was introduced into the luminal channel of blank microfluidic ECIS devices using a syringe pump at a flow rate of 1.3 mL/min (equivalent to a wall shear stress of 5 dyn/cm^2^). ECIS measurements were taken every 3 min before and after introduction of fluid flow for up to 50 minutes. Measurements in the blank devices with PBS indicated that ECIS resistance measurements were not affected by the application of fluid flow (**Figure 4A**), with no significant differences between the pre-flow and post-flow values for any of the electrode sizes at 4 kHz (**Figure 4B**). Further, a frequency scan showed impedance and resistance values were similar between pre-flow and post-flow conditions across frequencies (**Figure 4 C,D**).

**Figure 4:**
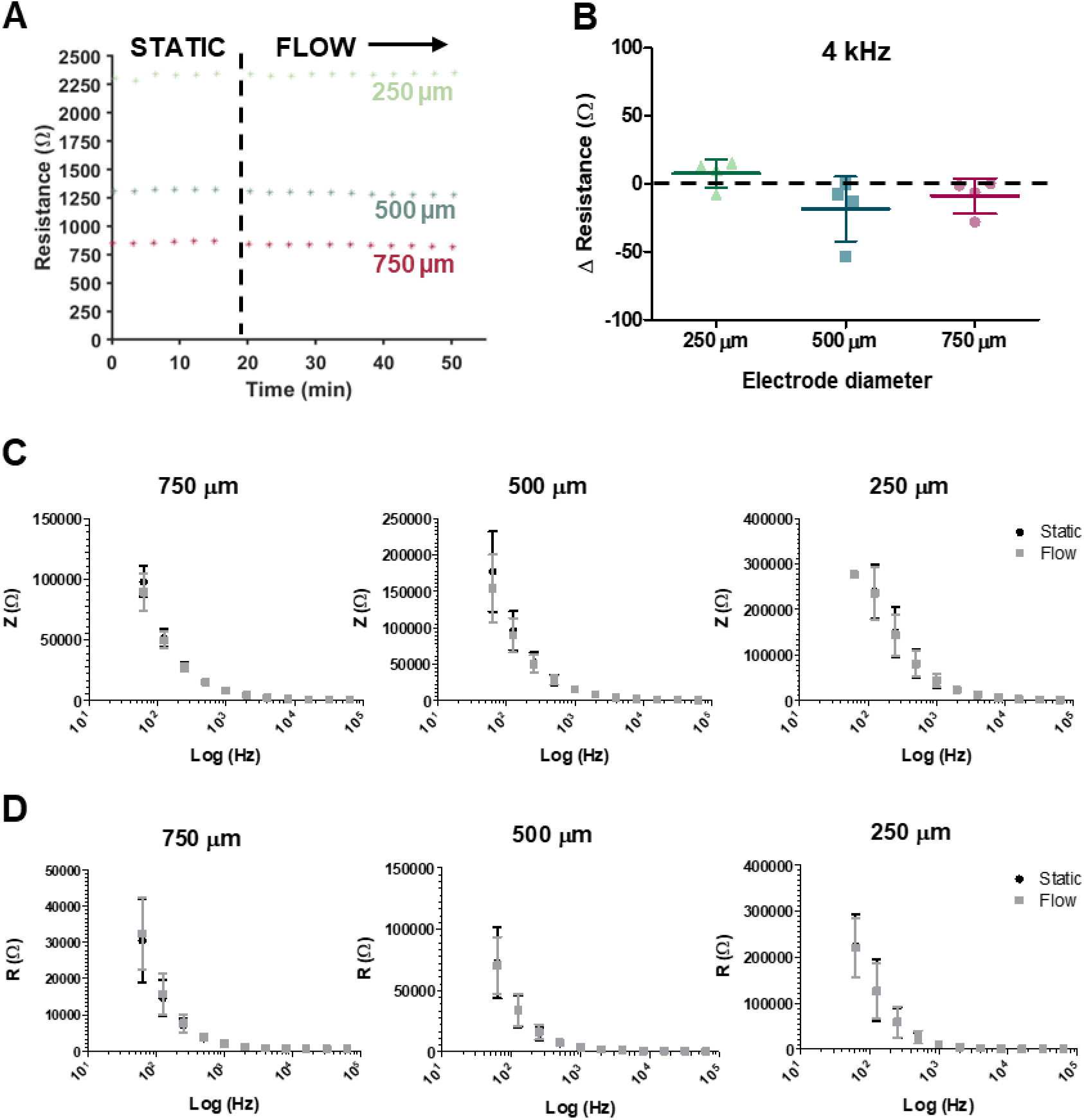
Microfluidic PM-ECIS measurements upon application of fluid flow. **A:** Resistance values measured at 4 kHz in blank devices filled with PBS before (static) and after application of fluid flow at 1.3 mL/min (equivalent to a wall shear stress of 5 dyn/cm^2^). **B:** No significant differences between pre-flow and post-flow values for any of the electrode sizes (Δ R = R_post-flow_ – R_pre-flow_). **C, D:** Frequency scan showing impedance values across scanned frequencies (**C**), and resistance (**D**). N=3-4 for each electrode size, data presented as mean ± standard deviation.

To test the platform’s capability for real-time ECIS measurement on perfused cultured cells, GFP-expressing HUVECs were seeded into the luminal channels, allowed to reach confluence over 2 days under static conditions, and then subjected to a perfusion flow rate of 20 µL/min. Parallel control cultures were maintained under static conditions. Live fluorescence imaging confirmed that cells perfused for up to 5 days remained viable and adhered to the porous membrane and electrodes comparably to cells grown under static conditions for the same duration (**Figure 5A**). Immunostaining for ZO-1 and VE-cadherin demonstrated that the perfused endothelium formed a confluent monolayer on the working (**Figure 5B**) and counter (**Figure 5C**) electrodes.

**Figure 5:**
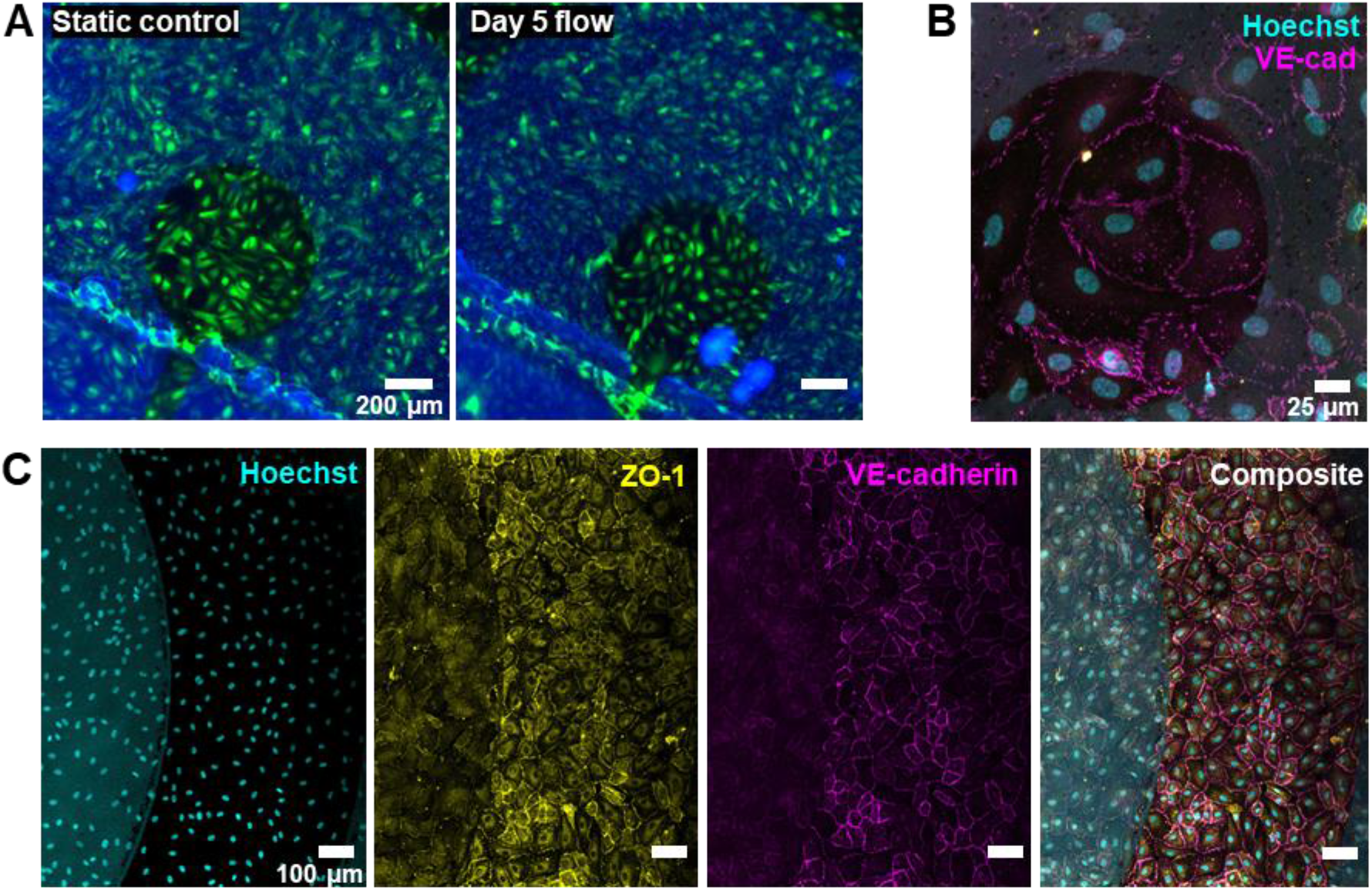
Confluent endothelial monolayer maintained after 5 days of perfusion culture in microfluidic PM-ECIS platform. **A:** Live fluorescence imaging of GFP-expressing HUVECs perfused for up to 5 days confirming viability and adherence of cells to porous membrane and electrodes vs. cells grown under static conditions for the same duration **B, C:** Immunostaining for ZO-1 and VE-cadherin and imaging with confocal microscopy confirms presence of confluent monolayer on working electrodes (**B**), and counter electrodes (**C**).

ECIS measurements were made continuously using three electrode sizes in each device prior to cell seeding, for 2 days post-seeding under static culture, and then for 3 days of either perfusion or static culture. In all cases, ECIS impedance (40 kHz) rapidly increased after cell seeding as the cells adhered and spread to form confluent monolayers, typically within one day of seeding (**Figure 6A,B,C**). ECIS resistance (4 kHz) also rose rapidly in this period but did not fully plateau after two days of static culture, suggesting cell junctions were continuing to mature at this time point (**Figure 6D,E,F**).

**Figure 6:**
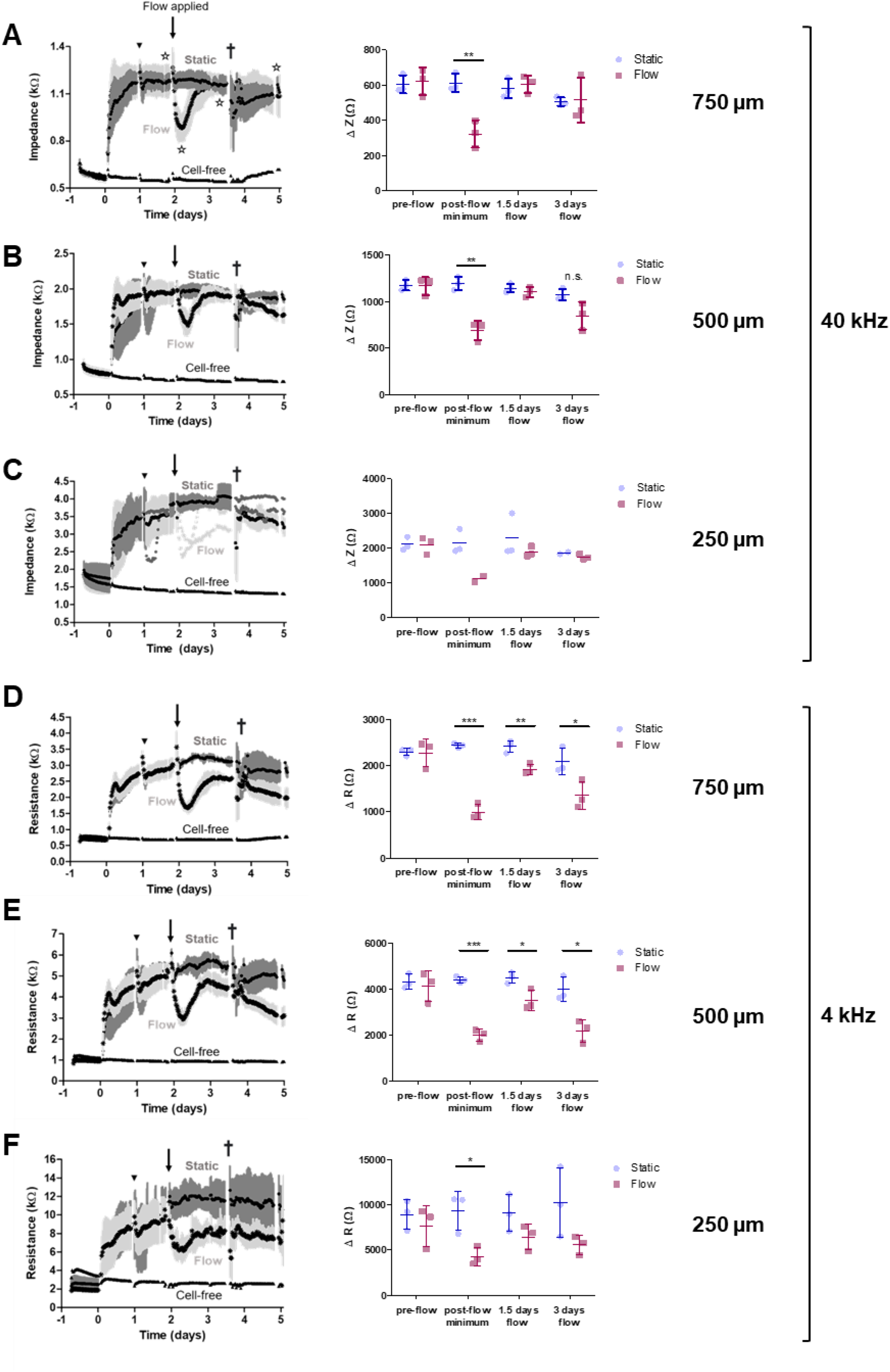
PM-ECIS measurements of HUVECs in perfusion culture. **A:** *Left:* Impedance measured at 40 kHz over 5 days of culture, 3 days of perfusion. Mean ± SD plotted for N=3 electrodes per condition, d = 750 µm. Stars indicate timepoints for impedance comparisons. *Right:* Comparison of impedance measured immediately before starting perfusion culture, minimum impedance value reached following start of perfusion, impedance after 1.5 days of flow, and 3 days of flow (Δ Z = Z_timepoint_ – Z_pre-cell seeding baseline_, similar calculation for resistance). **B:** Impedance at 40 kHz, d = 500 µm **C:** d = 250 µm; N=2-3 electrodes per condition. Mean ± SD plotted where applicable; for time intervals with N=2, individual replicates shown. **D, E, F:** Resistance measured at 4 kHz over 5 days of culture, 3 days of perfusion. Mean ± SD plotted for N=3 electrodes per condition, d = 750 µm (**D**), 500 µm (**E**), 250 µm (**F**). Cell seeding at t=0; black inverted triangle indicates manual media change on Day 1; † = syringes refilled with culture medium. Unpaired t-test, * = *p<0.05, ** = p<0.01, *** = p<0.001.* Data presented as mean ± standard deviation.

The initiation of perfusion at Day 2 resulted in significant decreases in impedance (**Figure 6A,B,C**) and resistance (**Figure 6D,E,F**) relative to static control cultures, with minimum values reached at 6.5 to 9.5 hours after induction of flow (p<0.01 for 750 and 500 µm electrode impedance at 40 kHz; p<0.05 for all electrode sizes for resistance at 4 kHz), indicating rapid flow-induced cell monolayer remodeling that decreased cell coverage and barrier integrity.

Within 24 hours of flow initiation, ECIS impedances in perfused cultures returned to levels similar to those of static control cultures **(Figure 6A,B,C**; p>0.3 for all electrode sizes on Day 1.5 after flow), indicating recovery of cell coverage on the ECIS electrodes and membranes. Impedances remained relatively constant for the remaining duration of the culture period, even when flow was paused briefly as the syringes were refilled with media at Day 3.5 (**Figure 6A,B,C**).

In contrast to impedance measurements, ECIS resistances measured at 4 kHz only partially recovered after flow initiation and trended lower over the 3 days of perfusion suggesting continued remodeling of the cell monolayer junctions over this period. This is corroborated by previous work from our group which shows increased electrochemical permeability in endothelium in the days following initiation of fluid flow^37^. In particular, resistance values measured by the larger 500 µm and 750 µm electrodes were significantly lower in the perfused cultures than in static both 1.5 days and 3 days after flow initiation (**Figure 6D,E**; p<0.05).

Similarly, resistances of perfused cultures measured by the smaller 250 µm electrodes trended lower than those of static cultures, but the differences were not significant (**Figure 6F**), in part because focal measurements with smaller electrodes result in great variability compared with more global measurements with larger electrodes^34^.

Nevertheless, the 250 µm electrodes were the most sensitive to changes in barrier integrity, both temporally and with respect to magnitude change in resistance. All three of the electrode sizes detected a significant decrease in resistance compared to static control at the post-flow minimum (**Figure 7A**), with the 250 µm electrodes showing the greatest magnitude decrease compared to 500 µm (p<0.01) and 750 µm (p<0.001). The 250 µm were also most sensitive to immediate change in barrier integrity following flow, with a significant decrease in resistance at 4 kHz (p=0.039) compared to static control measured within 30 minutes of applying fluid flow (**Figure 7B**); for the 500 and 750 µm electrodes this occurred at 1.5 hours post-flow (p=0.034 and p=0.026, respectively) (**Figure 7**, **Supplemental Table S1**). Change in impedance at 40 kHz occurred at 2 hrs (p=0.038) and 2.5 hrs (p=0.034) for 750 and 500 µm electrodes, respectively (**Supplemental Table S1**).

**Figure 7:**
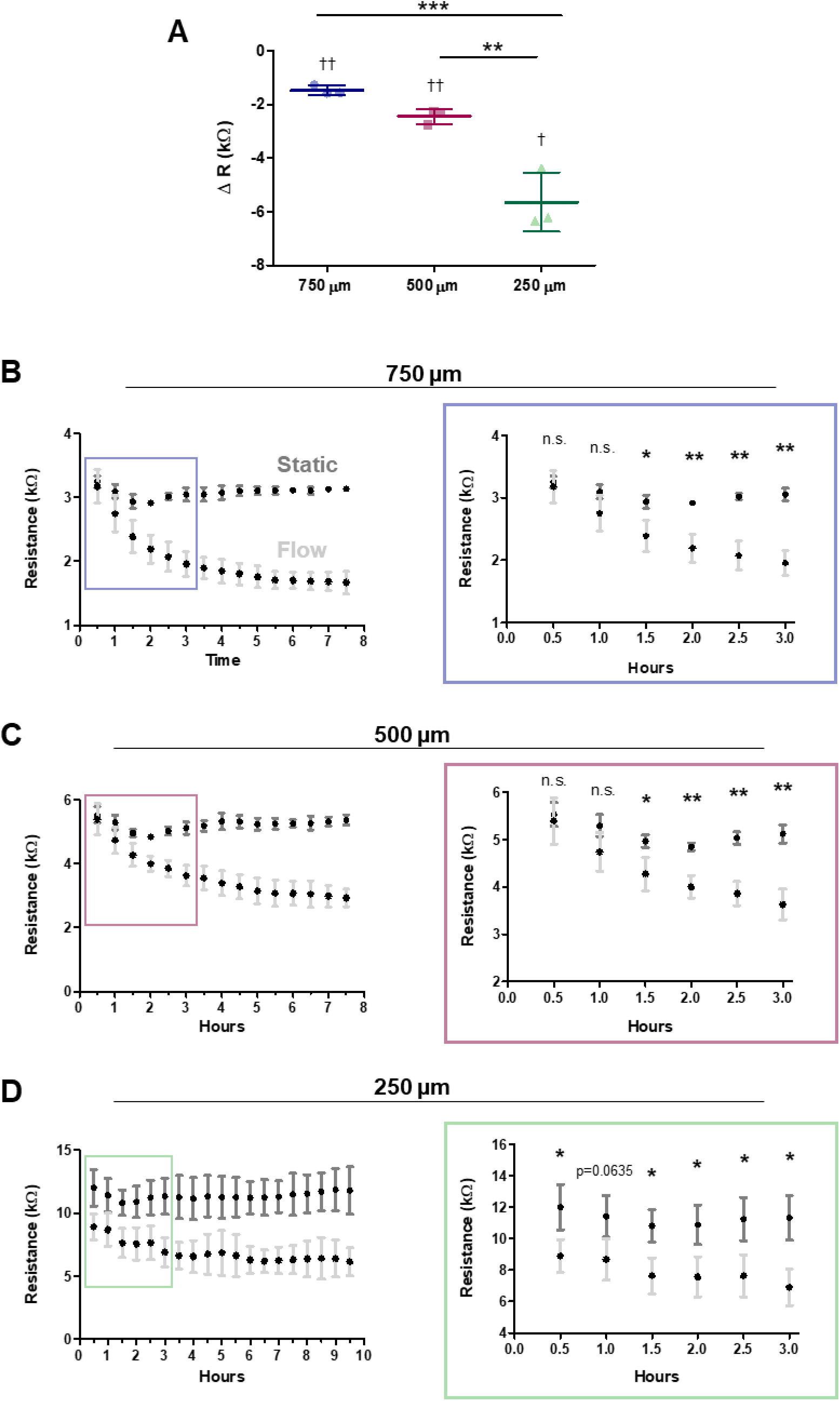
Electrode-size dependent response in resistance for HUVECs in perfusion culture. **A:** Decrease in resistance at 4 kHz after induction of flow (t=0) relative to static control. ΔR = R_flow_ – R_static_ for time point corresponding to post-flow minimum for each electrode. † p < 0.05, †† p<0.01 by one-sample t-test relative to change in ΔR=0; ** p < 0.01, *** p < 0.001 by one-way ANOVA with Bonferroni test. **B, C, D:** Resistance values for static and flow conditions over time up until post-flow minimum (*left*) and t = 3 hours (*right*) for 750 µm (**B**), 500 µm (**C**), 250 µm (**D**) electrodes. p < 0.05, ** p < 0.01 by unpaired t-test. N = 3 for each electrode size for each condition. Data presented as mean ± standard deviation.

## 4.0 Discussion

In this work, we addressed the pitfalls of the standard TEER measurement technique through a microfluidic embodiment of the PM-ECIS electrode platform. PM-ECIS was found to be robust to the presence or absence of hydrogel in the bottom channel, capable of continuous measurements over multiple days in co-culture and perfusion cultures, and sensitive to changes in cell coverage and barrier integrity induced by fluid flow.

### 4.1 Device fabrication

In this study, the device microfabrication method was selected with the goal of cost efficiency and ease of device assembly. The materials most commonly used in organ-on-chip devices - PDMS, silicon, glass, thermoplastics, 3D printing resins^38^ – often require time-consuming and costly fabrication techniques and materials. Double-sided adhesive has emerged as a cost-effective, rapid prototyping material for microfluidic device fabrication, which forgoes the technical challenges of assembly techniques required in typical fabrication such as chemical, plasma, heat or high pressure bonding^39^. Double-sided adhesive has been previously used for fabricating channels in microfluidic devices, though its application to OOC is still limited. In the last few years, characterization work performed on various biocompatible double-sided tapes has assessed properties like bonding strength, gas permeability and cytotoxicity^40,41^, as well as drug adsorption^42,43^. However, the majority of the publications for tape-based organ-on-chip systems have studied cells in monoculture^39–41,44–47^, with the exception of a hepatocyte-endothelium co-culture model designed by Xie *et al.*^48^. Evaluation of biological barrier formation and function in tape-based devices has also been limited – two studies have assessed epithelial barriers in multi-compartment devices^39,41^, and no studies thus far have done so with endothelial barriers. This is also the first incorporation of electrical sensors for real-time endothelial cell monitoring in a biocompatible tape-based OOC platform.

The integration of gold leaf PM-ECIS electrodes offers a high-fidelity, cost- and time-efficient alternative to the standard sputtering and e-beam metal deposition techniques. Moreover, the use of heat-bonded gold leaf enables a flat, continuous substrate with improved conductivity compared to sputter-coated membranes^33^. The successful implementation of PM-ECIS electrodes in an OOC platform is particularly relevant in the context of the recent movement toward developing simplified, lower-cost microfabrication processes^39,49^.

### 4.2 Collagen hydrogel ECIS resistance vs. TEER

Normalization of TEER values in culture setups containing hydrogel has been performed in cell culture inserts by subtracting resistance for hydrogel-containing inserts from the cell condition ^50,51^. Both Vila *et al.* and Singh *et al.* used chopstick electrodes and compared filter-only to filter + hydrogel resistance, finding there was no significant contribution by the hydrogel. Xu *et al.* implemented the same principle in a blood-brain barrier on chip model, subtracting the background resistance for collagen-only devices from BBB device resistance, without commenting on the magnitude of the collagen-only values^52^. Our own measurements using an equivalent electrode setup – electrodes positioned in the top channel inlet, and bottom “collagen” channel inlet – indicated a 1723 Ω contribution by the collagen gel (**Figure B**); using the standard TEER calculation, subtracting medium-only device resistance and normalizing to total exposed membrane area (1.9 cm^2^), this translates to a TEER value of 3274 ± 1416 Ω·cm^2^ for the collagen condition. Our study did not compare TEER values for a hydrogel device vs. cell-seeded device, therefore the degree to which the hydrogel TEER contribution would mask cell effects in our devices is unknown. However, Xu *et al.* measured a maximal TEER value of 1298 Ω·cm^2^ for BBB in flow conditions, compared to other conditions which had lower TEER: ∼600 Ω·cm^2^ for brain EC under flow, ∼400 Ω·cm^2^ for static BBB and ∼200 Ω·cm^2^ for static brain EC^52^.

The geometries of microfluidic devices exacerbate the non-homogenous current distribution inherent to this technique, with TEER measurements in microfluidic devices tending to be of greater magnitude than in cell culture inserts^7^. Even with the use of geometric correction factors^4,5^, however, simple subtraction of the hydrogel devices fails to take into account the cell distribution and behaviours that may induce changes in the hydrogel, as well as the inherent stability of the matrix^16^. Indeed, there is evidence to suggest that hydrogel contribution to impedance itself changes over time. Tonello *et al*., using a parallel electrode configuration, showed a decrease in impedance in cell-free gelatin-chitosan hydrogel over time across a range of frequencies (100 Hz – 10 kHz)^10^. The authors suggested this may be due to an increase in conductivity that occurs as the culture medium hydrates the gel; notably though, this effect was less pronounced in hydrogels embedded with stromal cells, which were presumed to act as insulators against the current. Conversely, De Leon *et al.*, using a planar electrode configuration, found that impedance of collagen gel increased with time over a range of 112 Hz to 31.2 kHz, which they attributed to protein and macromolecule deposition onto the electrodes^16^. The use of PM-ECIS, which measures cells directly grown on the electrodes, and is robust to the presence of hydrogel in other compartments (**Figure C**), circumvents these issues.

### 4.3 Blood-brain barrier model in microfluidic ECIS platform

Thus far there has been one example of ECIS electrodes incorporated onto porous membrane in a microfluidic BBB model – Matthiesen *et al*. used an interdigitated electrode configuration to monitor human induced pluripotent stem cell (hiPSC)-derived brain ECs and astrocytes, and the barrier’s functional response to N-acetyl cysteine amide^32^. Although their porous electrodes allow for more direct measurement of ECs that are in co-culture with astrocytes, the authors indicate that variability in metallization depth of the pores may contribute to greater error in ECIS measurements vs TEER. They also found that ECIS resistance for the BBB chips peaked at Day 1 vs. Day 2 for TEER values in the corresponding Transwell cultures, and that the metalized pores may have contributed to this difference.

In our microfluidic BBB culture model, the PM-ECIS resistance values (4 kHz) obtained for the 500 µm electrodes were similar to previous work from our group assessing co-culture of hCMEC/d3 + primary human astrocytes in PM-ECIS cell culture inserts^33^. The microfluidic BBB culture showed a comparable rate of increase and time to reach plateau in barrier integrity, with both microfluidic and cell culture insert platforms showing a difference of around 2.5 kΩ between co-culture and cell-free electrodes on the 3^rd^ day of culture. The peak resistance measured by the 250 µm electrodes (**Figure S1**) is also similar in magnitude to values obtained for brain microvascular endothelial cells cultured on solid-substrate ECIS^53^. Future studies could expand on this proof of concept model to test effect of co-culture and fluid shear on barrier integrity, as well as exploring other BBB-specific functional responses.

### 4.4 Microfluidic ECIS measurements upon application of fluid flow

The effect of long-term fluid flow (>24 h) on endothelial cell alignment and barrier integrity is well-established in the literature. However, there is also evidence that the effect of flow can become apparent at earlier timepoints. Both permeability to rhodamine-labeled albumin^54^ and forces across cell-cell junctions as measured by monolayer stress microscopy^55^ have been shown to increase within 1 hour of applying fluid shear to endothelial monolayers. ECIS electrodes have a high sensitivity for detecting cell movement and changes in endothelial barriers^20,56^, so we hypothesized that flow-induced monolayer changes could be detected by the platform on a short-term timescale.

To create a benchmark within our platform for the effect of flow in cell-free ECIS electrodes, PM-ECIS electrodes were submitted to 5 dyn/cm^2^ fluid shear, and shown to be robust to flow (**Figure 4**). The platform supported endothelial monolayer culture for up to 5 days of perfusion after 2 days in static culture, as confirmed by immunostaining for ZO-1 and VE-cadherin (**Figure 5**). Interestingly, this endpoint evaluation coincided with substantially decreased 40 kHz impedance and 4 kHz resistance values with respect to pre-flow monolayers. As an example, **Figure S2** displays the ECIS impedance and resistance up until the time of immunostaining, which ended at 66% and 53% of pre-flow values, respectively, for the 250 µm electrode shown in **Figure 5B**. This phenomenon is consistent with recent findings by Velasco *et al.*, which showed full electrode coverage by VE-cadherin-expressing HUVECs despite barrier resistance as low as 64% of pre-flow values^28^. This highlights the sensitivity and utility of PM-ECIS electrodes for providing additional information about the barrier. Our platform also produced a similar time course to Velasco *et al.* for decrease and partial recovery of endothelial resistance during fluid shear, with minimum resistance occurring 6-9 hours after the induction of flow in our devices (**Figures 6 and 7**) vs. 6 hours in the indicated study. The PM-ECIS electrodes were sensitive to these dynamic changes despite using very low fluid shear (0.06 dyn/cm^2^), with the 250 µm electrodes showing the greatest magnitude change in resistance in response to flow-induced changes in the endothelial barrier (**Figure 7**). This electrode size-dependent effect was consistent with the expected inverse proportional relationship between impedance and electrode size^22,57^, as well as with findings from our previous work^34^.

Interestingly, for the 250 µm electrodes, the induction of flow appeared to reduce the variability in resistance values in comparison to pre-flow values and to static condition (**Figure 7D**). This may be due to perfused cell culture getting a constant replenishment of nutrients, thereby maintaining more consistent cell health and reducing fluctuations in measurements. Alternatively, Velasco *et al*. propose that shear-induced cell reorientation may be less variable than the random individual cellular zigzag movements that occur in static monolayers, and that smaller electrodes would be more sensitive to this difference^28^. This study was limited to assessment of endothelial cells in perfusion culture; testing various fluid flow regimes could give a better understanding of these processes.

Shear stress is known to induce the reorganization of cell junctions and the cytoskeleton^58^, though the exact mechanisms through which this occurs are the subject of ongoing research. Endothelial barriers under flow show a decrease in TEER with fluid shear as low as 1 dyn/cm^2^ within 24 hours^59^. ECIS has been useful in providing a higher resolution picture of the time course of barrier response to flow. Studies with endothelial cells on solid substrate ECIS generally show an immediate increase in resistance, followed by a decrease that falls below the pre-flow value within a few hours^28,58,60^. The transient increase in resistance is presumed to be a response to the sudden onset of shear, and the corresponding magnitude and peak time is correlated with the magnitude of shear e.g. 5 dyn/cm^2^ leads to a 5% increase within 20 min, 50 dyn/cm^2^ to a 15% increase within 10 minutes^58^. Thus for a fluid shear of 0.06 dyn/cm^2^, as was used in this study, no appreciable transient peak in resistance would be expected. The apparent discrepancy between decreased ECIS values (**Figure S2**) and visually intact monolayer (**Figure 5**) is further explained by previous work which assessed expression of cell-cell junction markers by immunostaining, and showed no visible gap formation between endothelial cells despite using fluid shear conditions that induce significant changes in ECIS resistance^58^. Alternative analysis techniques can provide more insight into the specific processes that govern the endothelial response to flow. Steward *et al.* used monolayer traction microscopy and monolayer stress microscopy to generate high-resolution data on HUVECs subjected to fluid shear (10 dyn/cm^2^) at onset, 1 h, 12 h and 24 h of flow^55^. They found that laminar flow correlated to a 6% decrease in tractions exerted by the cells on the substrate over 24 h; in the same time period intercellular stresses decreased to less than half of pre-flow values. Furthermore, 1 h was enough time for the systematic alignment of intercellular stress to occur along the direction of fluid flow, with the effect becoming more pronounced by 12 h; conversely, alignment of the cell body only became apparent at 12 h. The application of of barrier disruptive agents like thrombin, or barrier protective agents like sphingosine-1-phosphate, produce changes in intercellular stress^55^; given that ECIS has a high detection sensitivity to transient responses to such stimuli^22^, these methods could potentially be used synergistically to assess flow response in endothelial barriers.

Analysis of the localization and the phenotype of cell-cell junctions may offer additional insight into ECIS measurements under flow. Ranadewa *et al.* showed that human cerebral microvascular endothelial cells exhibited a structural response to low fluid shear (1 dyn/cm^2^) over 24 h^61^. They found that laminar flow induced the relocalization of cell-cell junctions such as ZO-1 and VE-cadherin toward the cell center, and that this level of shear also corresponded to significantly smaller cell size compared to static control. Phenotype of cell-cell junctions – continuous, punctate, or perpendicular – has also been shown to play a role in endothelial barrier integrity. Gray *et al.* developed a tool called Junction Analyzer Program (JaNaP) which was used to investigate HBMEC monolayer response to barrier modulator cAMP, as well as show a difference between junction phenotypes in co-localization with local permeable regions^62^. Although global barrier assessment techniques (molecular permeability, TEER) were not shown to correlate with junction phenotype, the sensitivity of ECIS to local changes in the monolayer may make it amenable for being used in conjunction with such analysis approaches.

Although PM-ECIS resistance correlates with global barrier integrity measurements, including molecular tracer dye permeability^33^ and TEER^34^, ECIS is inherently a localized assessment of the cell layer. As such, the size and location of the electrodes in this study does not give a full picture about monolayer integrity throughout the channel. This can be addressed by increasing the relative area of electrode coverage through the channel, as has been done by Lewis *et al.* and Matthiesen *et al.*^27,32^, or incorporating multiple individual sensing electrodes across the length of the channel, as in the fluid flow array from Applied Biophysics^26^ to retain measurement sensitivity. Molecular permeability assays can be incorporated as well for further validation of barrier function.

One of the major challenges for incorporating TEER into organ on chip is that electrode size and placement is determined by device geometry, which impacts electric field uniformity; for example channel height has significant impact on the measured TEER values, as demonstrated by Yeste *et al*.^5^ Conversely, planar electrodes penetrate a small volume and distance^16^, which can be leveraged to measure just the cell layer of interest, and suggests that there would be less of an influence of device geometry on ECIS values. Indeed, this idea is supported by data from this study. The resistance values at 4 kHz for HUVECs in microfluidic PM-ECIS (Figure 6) closely match those produced by conventional solid substrate ECIS^20,63–66^, and this was reproducible across experimental runs (**Figure S3**). Moreover, they were comparable in magnitude to our previous work for the same HUVEC cell line grown in PM-ECIS cell culture inserts^34^. In the context of the significant challenges of translating TEER values between microfluidic and cell culture inserts, this data suggests the potential for PM-ECIS to act as a bridge between solid substrate ECIS, cell culture insert and membrane microfluidic barrier measurements.

## 5.0 Conclusions

In this study, we implemented gold leaf porous membrane ECIS in a microfluidic platform produced using cost-effective materials and fabrication techniques. A proof of concept, multi-day co-culture model of the BBB with inline electrical sensing was demonstrated in a tape-based organ-on-chip device. PM-ECIS electrodes were highly sensitive to flow-induced changes in a model of the endothelial barrier, in an electrode-size dependent manner. Our microfluidic PM-ECIS platform for the first time enables sensitive, non-invasive, real-time measurements of barrier function in OOC devices that incorporate critical physiological features like 3D co-culture, biomaterials and shear stress.

## CRediT authorship contribution statement

**Alisa Ugodnikov:** Conceptualization, Methodology, Formal analysis, Investigation, Data curation, Writing – Original draft, Project administration. **Joy Lu:** Investigation. **Oleg Chebotarev**: Methodology, Software, Funding acquisition. **Craig A. Simmons**: Conceptualization, Methodology, Resources, Writing – review, Supervision, Project administration, Funding acquisition.

## Appendix – Supporting Information

**Figure S1:**
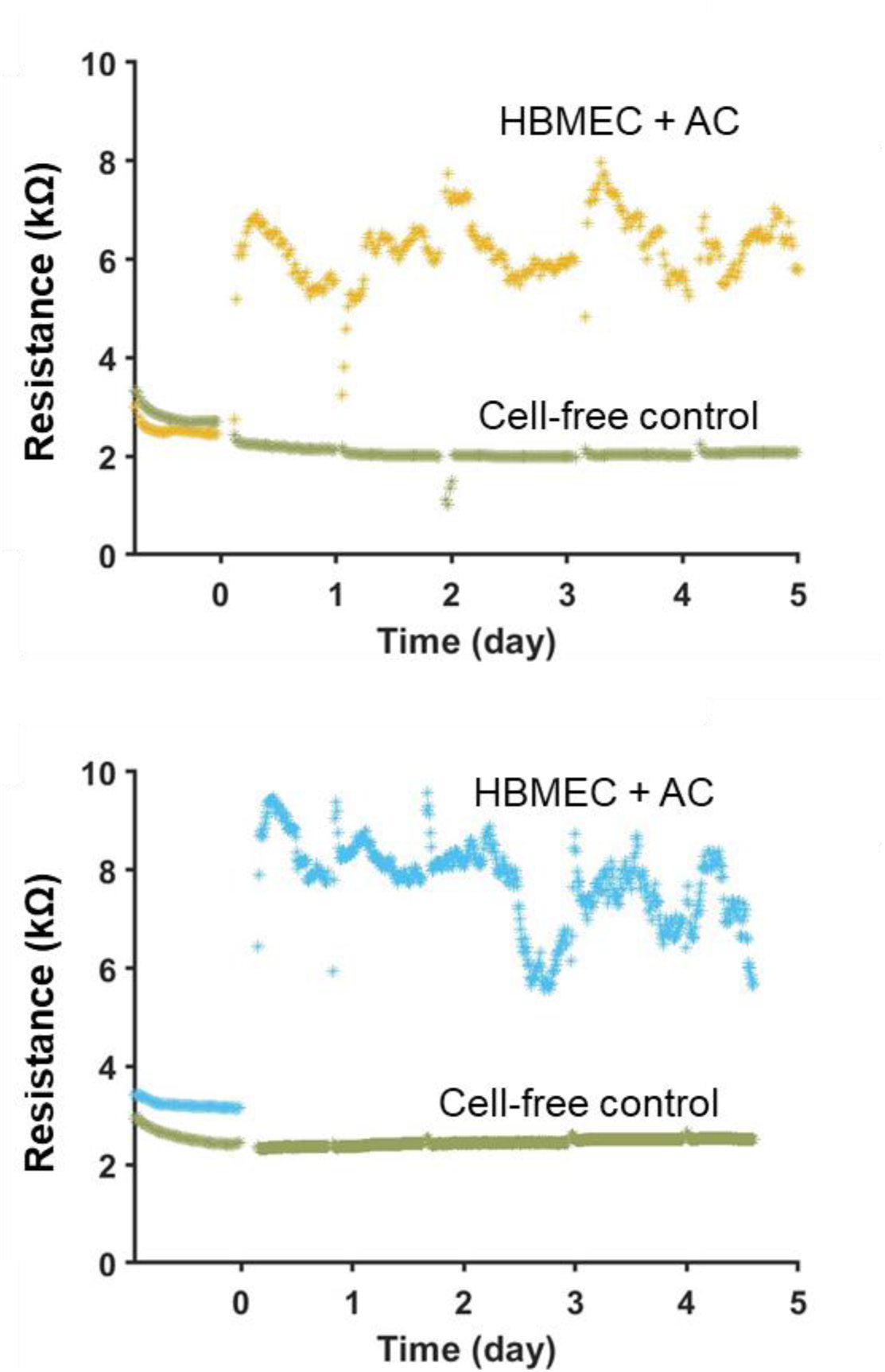
PM-ECIS resistance for microfluidic BBB co-culture model on 250 µm electrodes. Resistance at 4 kHz for HBMEC + primary human astrocyte co-culture vs. cell-free control, graphs showing example electrodes from different experiments.

**Figure S2:**
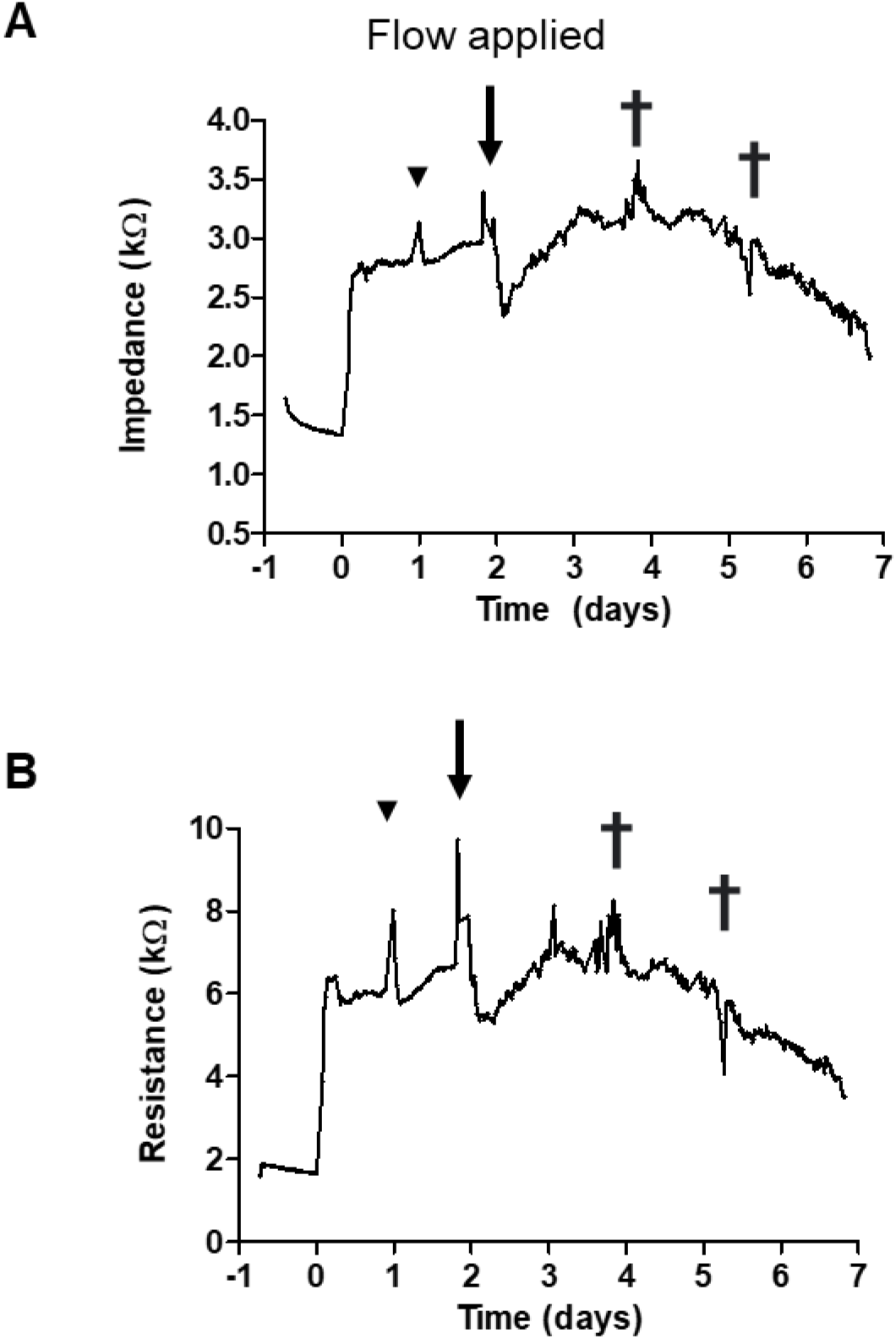
PM-ECIS measurements of HUVECs in perfusion culture. **A:** Impedance measured at 40 kHz and **B:** resistance measured at 4 kHz over 7 days of culture, 5 days of perfusion, corresponding to 250 µm working electrode shown in Fig. 5B. Cell seeding at t=0; black inverted triangle indicates manual media change on Day 1; † = syringes refilled with culture medium.

**Figure S3:**
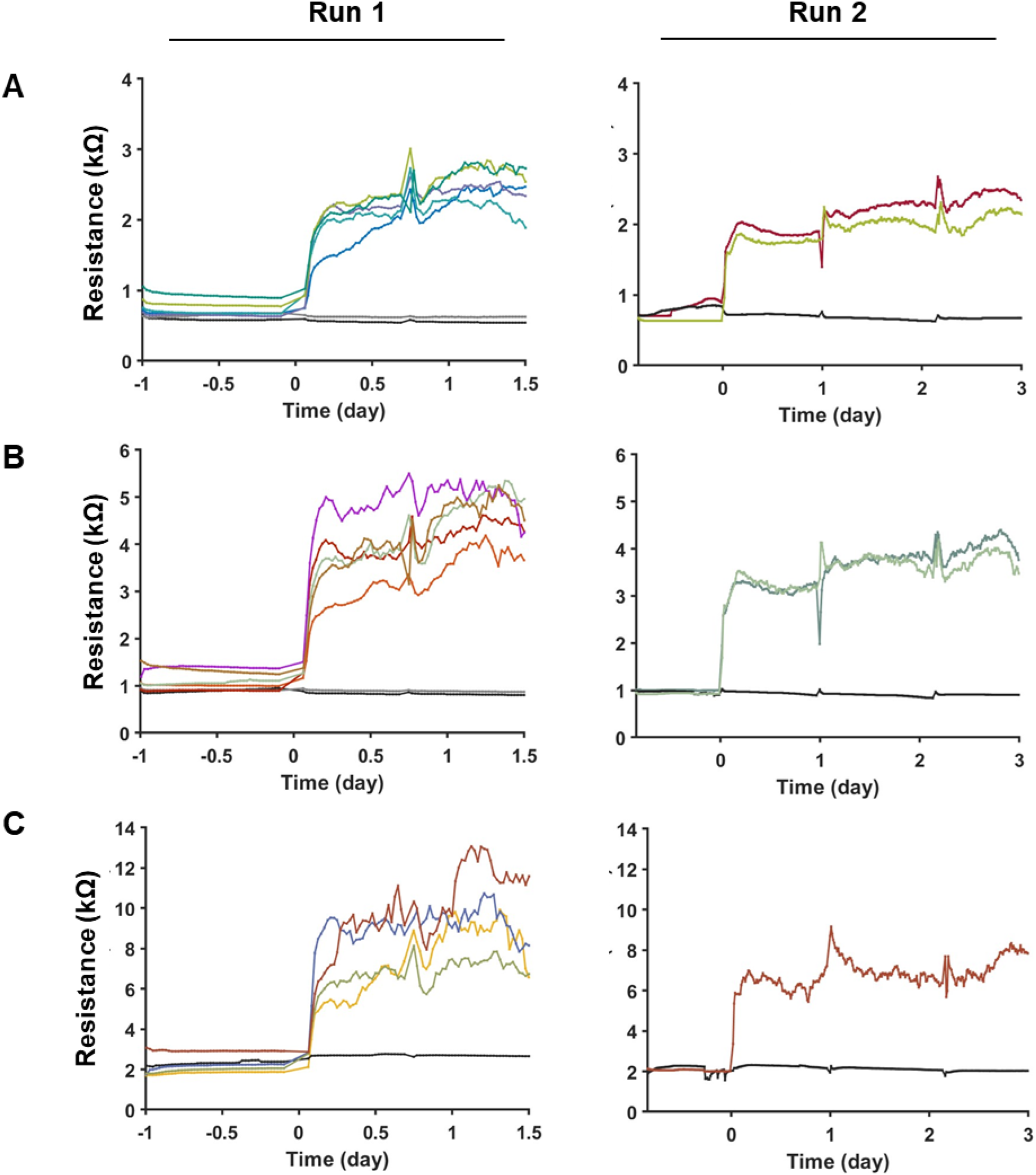
Microfluidic PM-ECIS measurements of HUVECs for trials run of static culture. Resistance at 4 kHz measured over time for **A**: 750 µm, **B**: 500 µm, and **C**: 250 µm working electrodes. Cell seeding at t=0.

**Table S1:** Time after induction of flow for significant change in PM-ECIS values for HUVECs in perfusion.

|  | Impedance at 40 kHz |  | Resistance at 4 kHz |  |  |
| --- | --- | --- | --- | --- | --- |
| Time (hours) | 750 $\mu\text{m}$ | 500 $\mu\text{m}$ | 750 $\mu\text{m}$ | 500 $\mu\text{m}$ | 250 $\mu\text{m}$ |
| **0.5** | n.s.<br>(p=0.5667) | n.s.<br>(p=0.2073) | n.s.<br>(p=0.6806) | n.s.<br>(p=0.6871) | *<br>**(p=0.039)** |
| **1** | n.s.<br>(p=0.4051) | n.s.<br>(p=0.2707) | n.s.<br>(p=0.1185) | n.s.<br>(p=0.1097) | *n.s.*<br>(p=0.0635) |
| **1.5** | n.s.<br>(p=0.1099) | n.s.<br>(p=0.1592) | *<br>**(p=0.0257)** | *<br>**(p=0.0343)** | *<br>**(p=0.0229)** |
| **2** | *<br>**(p=0.0396)** | *n.s.*<br>(p=0.0715) | **<br>**(p=0.0047)** | **<br>**(p=0.0042)** | *<br>**(p=0.0321)** |
| **2.5** | *<br>**(p=0.0239)** | *<br>**(p=0.0337)** |  |  |  |
| **3** |  | *<br>**(p=0.026)** |  |  |  |

